# Working Memory and Impulsivity and Artificial Neural Networks

**DOI:** 10.1101/2020.10.26.355990

**Authors:** Markus Ville Tiitto, Robert A. Lodder

## Abstract

Attention deficit hyperactivity disorder (ADHD) is a neurodevelopmental disorder characterized by inattention, hyperactivity, and impulsivity. The treatment of ADHD could potentially be improved with the development of combination therapies targeting multiple systems. Both the number of children diagnosed with ADHD and the use of stimulant medications for its treatment have been rising in recent years, and concern about side-effects and future problems that medication may cause have been increasing. An alternative treatment strategy for ADHD attracting wide interest is the targeting of neuropsychological functioning, such as executive function impairments. Computerized training programs (including video games) have drawn interest as a tool to train improvements in executive function deficits in children with ADHD. Our lab is currently conducting a pilot study to assess the effects of the online game Minecraft as a therapeutic video game (TVG) to train executive function deficits in children with ADHD. The effect of the TVG intervention in combination with stimulants is being investigated to develop a drug-device combination therapy that can address both the dopaminergic dysfunction and executive function deficits present in ADHD. Although the results of this study will be used to guide the clinical development process, additional guidance for the optimization of the executive function training activities will be provided by a computational model of executive functions built with artificial neural networks (ANNs). This model uses ANNs to complete virtual tasks resembling the executive function training activities that the study subjects practice in the Minecraft world, and the schedule of virtual tasks that result in maximum improvements in ANN performance on these tasks will be investigated as a method to inform the selection of training regimens in future clinical studies.

## Background

Attention deficit hyperactivity disorder (ADHD) is a neurodevelopmental disorder characterized by inattention, hyperactivity, and impulsivity^1^. The onset of ADHD typically occurs by 3 years of age, and must occur by 12 years of age for a diagnosis. While symptom severity decreases with age, ADHD may persist into adulthood^2^, when the hyperactive-impulsive symptoms typically subside but the inattentive symptoms persist^3^. The cause of ADHD remains largely unknown, but dopaminergic dysfunction has been implicated as playing an important role. For example, stimulant medications (such as methylphenidate and amphetamine) that increase dopaminergic neurotransmission are the most efficacious treatment for ADHD, and investigations into genetic factors of ADHD have revealed modest associations for the dopamine transporter (DAT1) and dopamine receptors D4 & D5 (DRD4, DRD5)^4^. However, ADHD has a considerably high estimated heritability rate^5^, so these modest associations observed for dopaminergic system genes indicate that many other factors also play an important role. Therefore, the treatment of ADHD could potentially be improved with the development of combination therapies targeting multiple systems.

Both the number of children diagnosed with ADHD and the use of stimulant medications for its treatment have been rising in recent years^6^. The overall prevalence of ADHD increased from 6.1% in 1997-1998 to 10.2% in 2015-2016. The prevalence in boys increased from 9.0% to 14.0%, while the prevalence in girls more than doubled from 3.1% to 6.3%. However, it is unknown at this time whether these increases reflect the actual number of ADHD cases, or are instead a result of factors leading to increased diagnosis of ADHD, such as increased physician awareness, changes in diagnostic criteria, or increased access to medical care. In 2011, 3.5 million children were being treated with stimulants according to parents^7^. In addition, prescriptions for ADHD medications in women of child-bearing age increased by 344% from 2003 to 2015, which included a 700% increase in women of ages 25-29 years and a 560% increase in women of age 30-34 years^8^.

Despite the variety of pharmacologic treatment options available and their widespread use, there still remains a strong need to develop additional therapies^9^. While a general consensus exists for the efficacy of stimulants (the first-line therapy for ADHD)^10^, a recent systematic review of methylphenidate concluded that the evidence supporting its use is of very low-quality, and more caution should be exercised in its use^11^. In addition, the benefits that result from stimulant use do not persist after discontinuation and patients with ADHD still suffer from adverse long-term outcomes such as poor academic performance, drug addiction, and criminal behavior to a much greater degree than non-ADHD subjects despite optimal therapy^12^. Despite their tolerability in the majority of ADHD patients, the side effect profile of stimulants (anxiety, irritability, insomnia, gastrointestinal distress, loss of appetite, and growth suppression) still precludes their use in a significant number of patients due to the widespread prevalence of their prescribing. An additional safety concern of stimulants are their long-term effects, for which there is a paucity of research in humans. However, recent studies have shown altered cerebral blood flow responses after discontinuation of stimulant use^13^, reduced GABA levels in the pre-frontal cortex of ADHD patients treated with stimulants at a young age^14^, and altered white matter in children treated with methylphenidate^15^. Drug treatments also suffer from further under-utilization due to parents’ concerns about their safety and preference for the use of non-drug treatments^9^. Finally, the widespread use of stimulants has also led to their misuse and abuse for non-medical purposes, which may be obviated by a greater availability of other effective treatments for ADHD.

An alternative treatment strategy for ADHD attracting wide interest is the targeting of neuropsychological functioning, such as executive function impairments. Russell A. Barkley proposed an executive function theory for ADHD in 1997, which states that an impairment in the core executive function inhibition is the central causative factor in the development of ADHD^16^. Since the hyperactive behavior of children with ADHD reflects a lack of behavioral inhibition and the use of inhibition keeps attentional resources available for the use of other executive functions, Barkley reasoned that a deficit in inhibition leads to a cascade of impairments in other executive functions culminating in the characteristic behavior of ADHD.

Executive functions are a set of effortful, top-down mental processes that govern attention and regulate behavior^17^. Executive functions enable both visualization of the future and remembrance of the past, allowing for control of one’s behavior over time to accomplish long-term goals and self-reflection to recognize past mistakes so that they are not repeated. In addition to the consideration of behavior across time, executive functions also enable conscious manipulation of thoughts and ideas, which includes the use of creativity to combine conflicting ideas in novel ways.

A set of core executive functions has been proposed to serve as a foundation for higher order executive functions and the general application of executive functioning in life activities^17^. Application of the executive functions allows for the development and use of important skills such as time management, organization, planning, self-regulation, sustained attention, and metacognition. Two examples of core executive functions are working memory and inhibition, which work together very closely. Working memory is the ability to hold a piece of information in consciousness (short-term memory) and then manipulate it in some way^18^. Inhibitory control includes the ability to ignore environmental distractions (attentional control)^19^ as well as the ability to prevent automatic or impulsive thoughts and behaviors when they may not be appropriate.

Computerized training programs (including video games) have drawn interest as a tool to train improvements in executive function deficits in children with ADHD^19–21^. Our lab is currently conducting a pilot study to assess the effects of the online game Minecraft as a therapeutic video game (TVG) to train executive function deficits in children with ADHD^22^. The effects of the TVG intervention in combination with stimulants is being investigated to develop a drug-device combination therapy that can address both the dopaminergic dysfunction and executive function deficits present in ADHD. Although the results of this study will be used to guide the clinical development process, additional guidance for the optimization of the executive function training activities will be provided by a computational model of executive functions built with artificial neural networks (ANNs)^23^. This model uses ANNs to complete virtual tasks resembling the executive function training activities that the study subjects practice in the Minecraft world, and the schedule of virtual tasks that result in maximum improvements in ANN performance on these tasks will be investigated as a method to inform the selection of training regimens in future clinical studies.

Artificial neural networks (ANNs) are a group of computational models that are inspired by the structure and function of biological neural networks^23^, and are part of a broader collection of computational techniques called machine learning algorithms, which are a set of computational models that learn how to complete tasks more accurately by performing the tasks on their own without explicit guidance from a human^24^. ANNs are composed of individual units that receive inputs, process these inputs, and produce an output, similar to the way that individual neurons in biological neural networks receive, process, and produce electrical signals. Multiple units are combined into layers, and these layers are combined to form the full network. Each unit in a given layer receives multiple inputs produced by units in the previous layer, and produces a single output that is used as an input for multiple units in the next layer. The design of the units, layers, and their connectivity pattern is known as the network architecture.

ANNs have attracted interest as a computational model in drug development and healthcare because of their ability to learn how to accomplish tasks that involve the processing of complex input data. For example, ANNs have been used to predict the quantitative structure-activity relationships of potential drug candidates^25^, generate novel chemical structures for drug candidates^26^, predict the occurrence of cardiovascular disease^27^, and screen for skin cancer^28^, diabetic retinopathy^29^, and other retinal diseases^30^. This automation of complex tasks requiring specialized knowledge may offer a significant potential advantage in time and cost savings for the healthcare system.

Due to their similarity to biological neural networks, ANNs could potentially be used as a computational model for neurological activity. This premise inspired the selection of ANNs as a computational tool to simulate the training of executive functions in this project. Importantly, the neurological activities of interest in this project are the neural adaptations that occur during the learning process, rather than the actual neurological activity that occurs during the use of executive functions. While it is recognized that ANNs are extensively simplified approximations for biological neural activity, they are nonetheless being developed to perform complex human tasks such as goal-oriented conversations^31–32^ and navigating autonomous vehicles^33^. Thus, it is hypothesized that ANNs can be trained to perform virtual tasks that resemble activities in humans that require the use of executive functions, and that this training process in ANNs can provide insight into the optimal way to train the use of executive functions in humans.

The construction of a computational model for executive function training would enable the rapid *in silico* simulation of different combinations and schedules of these activities (Figure 1). If a virtual set of activities can be designed to resemble the use of human executive functions closely enough, these simulations could potentially provide insight into how variations in the selection and scheduling of these activities affect the outcomes of executive function training in humans. An examination of the effects of varying combinations of these virtual activities on ANN performance would inform the selection of an optimal schedule of activities to improve executive function deficits in humans. In this way, an effective virtual simulation model of executive function training activities could provide considerable time and cost savings compared to clinical studies for the optimization of the executive function training activity schedules in humans.

**Figure 1.**
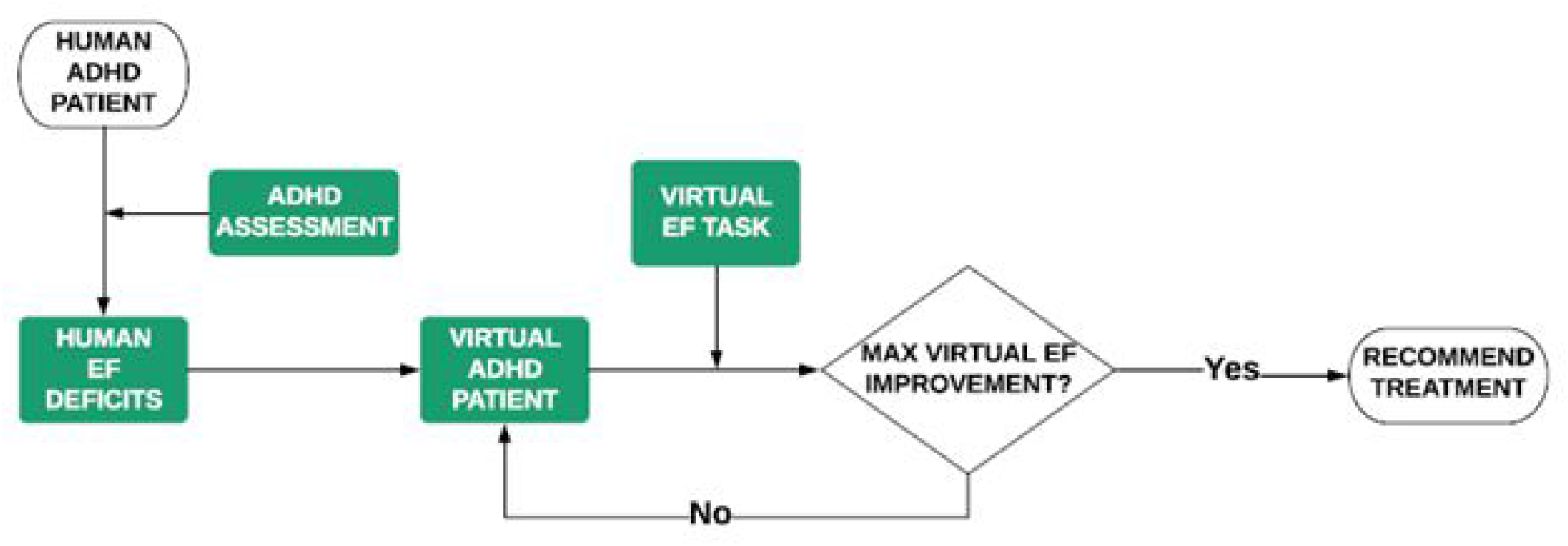
Generation of Personalized Executive Function Training Schedules

The TVG plus stimulants drug-device combination therapy will utilize a novel personalized medicine approach where an individualized treatment regimen consisting of an initial stimulant dose recommendation and schedule of therapeutic video game activities will be determined from the initial ADHD assessment results of new patients. In the computational executive function model, a set of ANNs are used to represent a virtual “subject” comprised of a set of executive functions that work together to perform the virtual executive function tasks. As a virtual “subject” completes virtual executive function tasks, the ANNs representing its executive functions will undergo further training and the “subject’s” performance on these tasks will improve. Multiple combinations and schedules of virtual executive function tasks can then be simulated rapidly in each virtual “subject” to determine an optimal training regimen to target the pattern of executive function deficits in an individual patient.

The objectives of this work are to create an ANN-based representation of the core executive function working memory, create groups of virtual “subjects” differentiated by the performance of this working memory representation, and create an impulsivity function that can generate automatic behaviors that do not result from the use of executive functions. The impulsivity function was created as an initial step towards the development of a representation for the core executive function of inhibition, and was inspired by the race model^34^ and passive dissipation hypothesis^35^ for behavioral inhibition. This work will serve as an initial starting point for the later addition of more executive function representations, and the creation of virtual executive function tasks that require the use of the executive function representations for their completion.

## Methods and Results

### Generating Working Memory Deficiencies (Colab Notebook)

As a first step towards the creation of a computational model of working memory deficiencies, convolutional neural networks (CNNs) were trained to identify handwritten digits in the MNIST dataset. The Keras Machine Learning library^36^ was used to create and train the CNNs and the experiment was run using Python v3 in a Google Colaboratory Notebook. In this implementation, the input of an MNIST image file into the first layer of the network is proposed to represent the holding of information in consciousness component of working memory. The subsequent processing of this image pixel data by the internal network layers is proposed to represent the information manipulation component of working memory. Two groups of CNNs were created by varying the number of MNIST images (the size of the training set) used in their training process. A “healthy” control working memory group consisted of CNNs trained to achieve high accuracy in the handwritten digit recognition task, while a “deficient” working memory group consisted of CNNs trained to achieve approximately half the accuracy of the “healthy” control group.

All of the CNNs in both groups possessed an identical architecture, or structure of layers and connectivity between individual units. The architecture chosen was a modified version of the original historic CNN called lenet that was developed to identify handwritten digits in MNIST to automate zip code recognition for postal service^37–38^. The CNN architecture used here consists of two sets of convolutional and pooling layers, followed by two fully-connected layers, and a softmax classifier. The first and second convolutional layers consisted of 20 filters and 50 filters, respectively, each with a 5×5 kernel and rectified linear unit (ReLU) activation function^39^. Both pooling layers used a 2×2 filter with stride = 2. The first fully-connected layer contained 500 units with the ReLU activation function, and was followed by a 10-unit fully-connected layer with the softmax activation function. A batch normalization layer^40^ was included after each convolutional layer and the first fully-connected layer before the ReLU activation function. The maximum value of the softmax activations was selected as the final output.

To generate deficiencies in this working memory representation, the relationship between the quantity of training examples and handwritten digit recognition performance of CNNs was first investigated. Five groups of identically structured CNNs (n = 10 x CNNS per group) were trained with sets of MNIST images with sizes in the range of 25 – 50000 MNIST images per set. The MNIST images used for training were randomly selected from the training subset of the MNIST dataset, and randomly selected sets containing less than 100 MNIST images were checked to ensure that they contained at least one example image of each handwritten digit 0-9. Updates of the CNN parameters (weights & biases) were performed after training on batches of 25 MNIST images. The categorical cross-entropy loss function^41^ was used to measure the handwritten digit recognition performance of the CNNs during training and calculate the gradients of its weights and biases. The ADAM optimizer^42^ was then used to calculate the magnitude of the parameter updates from these gradients.

After training, the percent accuracy of handwritten digit recognition performance of each CNN was evaluated on the 10,000 MNIST images of the MNIST test set, and the mean accuracy of each group of CNNs was determined. As expected, the group mean handwritten digit recognition accuracy increased with increasing number of MNIST images presented during training (Figure 2). The minimum group accuracy achieved was 44.3% (SD 4.1%) with 25 MNIST training images, and the maximum group accuracy achieved was 98.2% (SD 0.5%) with 50,000 MNIST training images. The observed relationship between group accuracy and the number of MNIST training images was logarithmic, and the rate of increase in performance was relatively minimal in groups trained with set sizes larger than 2,500 MNIST images (mean group accuracy = 95.7% (SD 0.5%)).

**Figure 2.**
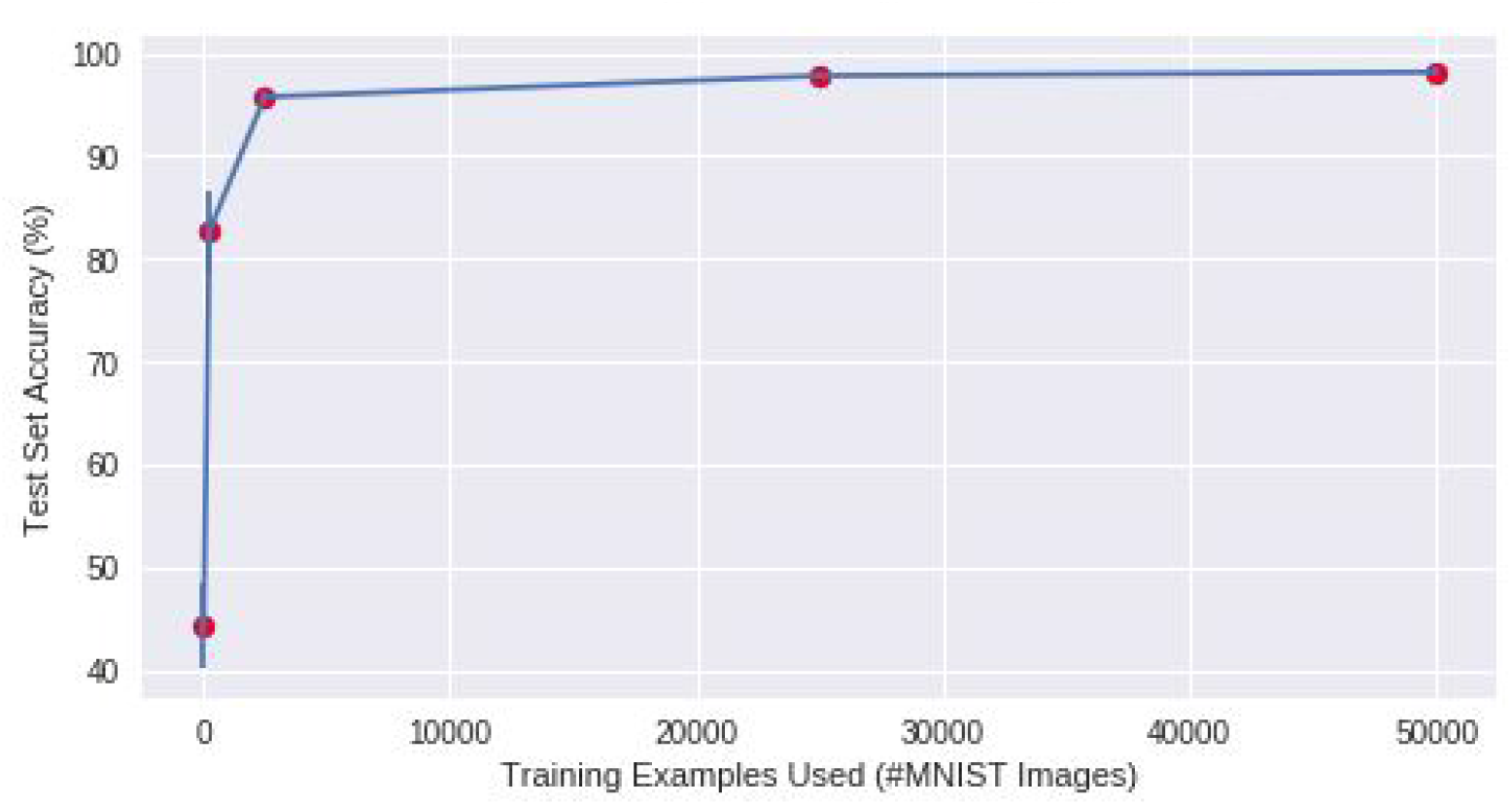
Effect of MNIST Training Set Size on MNIST TestSet Handwritten Digit Recognition of CNNs (n = 10 x CNNs per Group)

Training procedures for the CNNs representing working memory were selected to create a sizeable difference in performance between the “healthy” working memory control group with high handwritten digit recognition performance and the “deficient” working memory group with poor handwritten digit recognition performance in a minimal amount of computational time. Computational efficiency was prioritized over marginal improvements in accuracy in the selection of a training procedure for the “healthy” working memory control group. While a 2.5% improvement in group accuracy was observed from increasing the training set size from 2,500 to 50,000 MNIST images, this improvement was achieved at a significant cost of increased computational time. Thus, a training set size of 2,500 MNIST images was selected to efficiently train CNNs in the “healthy” working memory control group to achieve a sufficiently high handwritten digit recognition performance.

The selection of a training procedure for the “deficient” working memory group was based on a similar consideration of weighing the level of performance achieved with the training time required to achieve the performance. A large performance decrement compared to the “healthy” working memory group was desirable in this case. The minimum accuracy achieved in this experiment was 44.3% (SD 41%) in the group trained with 25 MNIST images. While this level of accuracy could be decreased further by reducing the size of the training set, this decrease was achieved at a cost of significantly increased computational time to randomly select smaller training sets with at least one MNIST image containing each handwritten digit. Thus, a training set size of 25 MNIST images was selected to efficiently train CNNs in the “deficient” working memory group to achieve a sizable reduction in handwritten digit recognition performance compared to the “healthy” working memory control group.

To summarize, this investigation was performed to select training procedures for MNIST handwritten digit recognition by CNNs as a computational representation for working memory in humans. In this context, a “healthy” working memory was defined as high handwritten digit recognition performance by CNNs and a “deficient’ working memory was defined as poor handwritten digit recognition performance by CNNs. Differences in performance were created by varying the number of MNIST images used to train the CNNs. A training set size of 2,500 MNIST images was selected to generate CNNs with high handwritten digit recognition performance representing “healthy” working memory, while a training set size of 25 MNIST images was selected to generate CNNs with poor handwritten digit recognition performance representing “deficient” working memory.

### Prepotent Impulsivity

As a first step towards the construction of a computational model for the behavioral inhibition executive function, the prepotent impulsivity function was created to generate impulsive behaviors that may be inhibited. Prepotent responses are defined as unproductive behaviors that have been overlearned in a given circumstance, and are then used indiscriminately in other circumstances where they may no longer be appropriate^16,43^. This function was designed to be an addition to the MNIST handwritten digit recognition representation of working memory and activates unproductive behaviors resembling prepotent responses in this context.

When the convolutional neural networks representing working memory were evaluated for their performance, their handwritten digit recognition accuracy was determined on an ordered test set. In other words, the convolutional neural networks were first presented with all the images containing a handwritten zero, then all the images containing a handwritten one, all the images containing a handwritten two, and so on. In contrast, this experiment was conducted with shuffled test sets where the images containing the various handwritten digits were presented to the convolutional neural networks in a random order. A shuffled test set will contain randomly distributed sub-sequences of consecutive, but different, images containing the same handwritten digit. When these sub-sequences with consecutive images containing the same handwritten digit are presented to a convolutional neural network, a prepotent impulse is created. This prepotent impulse grows in strength as the number of consecutive digit repeats grows larger and promotes an automatic response with the identity of the repeated digit from the virtual “subject”. The automatic prepotent response is produced without the presentation of an MNIST image to the convolutional neural network, and is more likely to be incorrect than a non-impulsive response produced with the presentation of an MNIST image to the convolutional neural network. Thus, the prepotent impulsivity function can be considered a method to produce erratic behaviors without the benefit of reasoning with executive functions (working memory) in this model.

In addition to the prepotent impulse, the prepotent impulsivity function also includes a competing executive function-activating component. The executive function-activating component supports the activation of a response determined by executive functions, specifically the presentation of an MNIST image to the convolutional neural network representing working memory. The probability of carrying out an impulsive prepotent response is determined by subtracting the strength of this executive function-activating component from the strength of the prepotent impulse (Figure 3).

**Figure 3.**
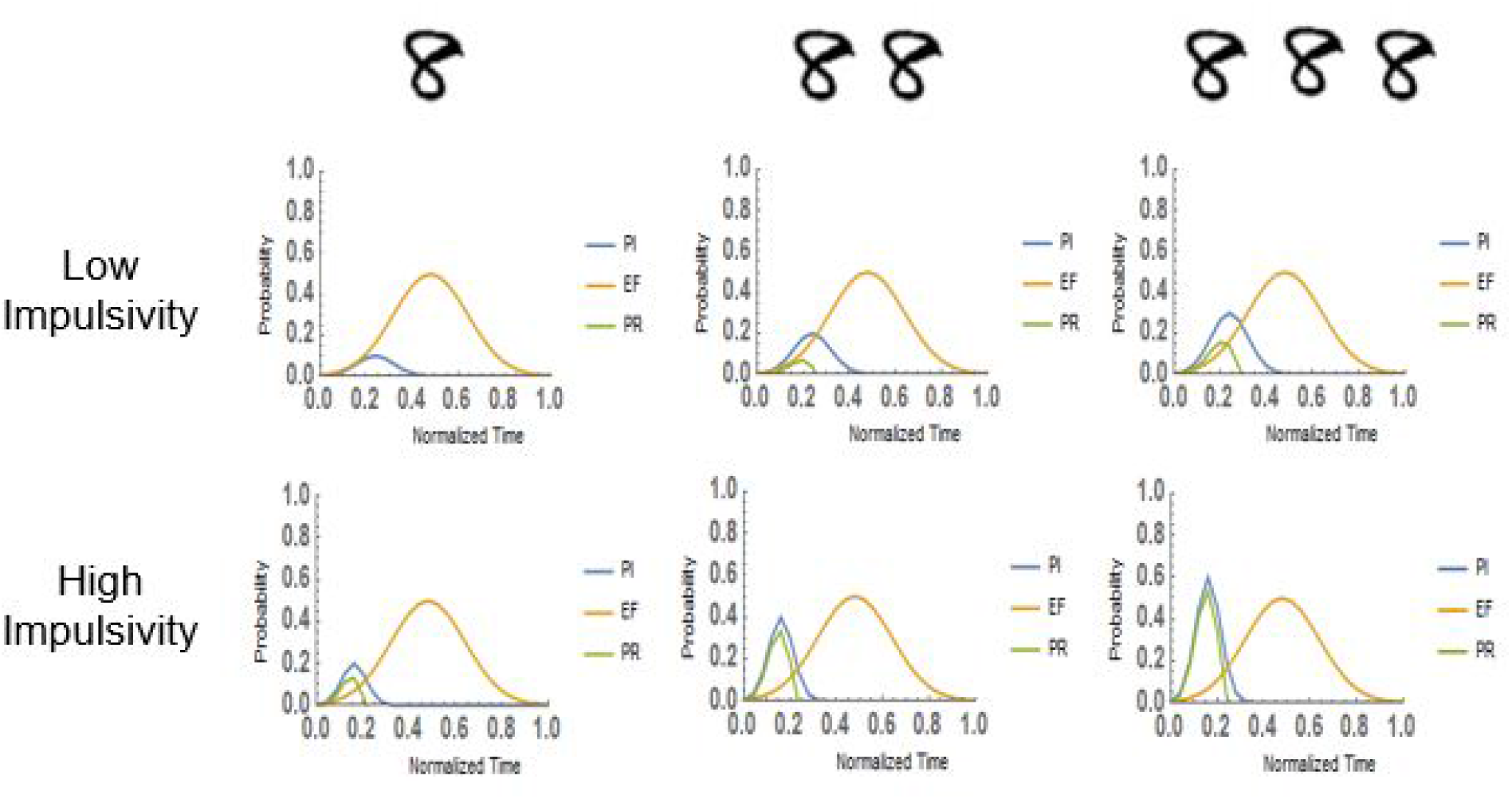
Effects of Repeated Handwritten Digits on Prepotent Impulsivity Function (PI = Prepotent Impulse; EF = Executive Function-Activating Component; PR = Prepotent Response Probability)

Both the prepotent impulse (PI curve) and the executive function-activating component (EF curve) of the prepotent impulsivity function are Gaussian functions calculated with time as the independent variable. This time value is considered an abstract representation of computational processing time and not a real time related to the computational task. Both curves are constructed with a strength parameter (*α*), an efficiency parameter (*β*), and a delay parameter (*γ*).

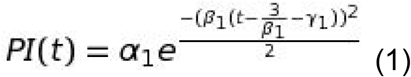

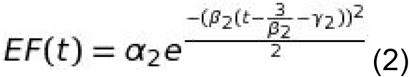

The strength parameter determines the magnitude of the curve’s peak, the efficiency parameter determines the rate at which the curve reaches its peak, and the delay parameter determines the location where the curve begins to grow. While the values of the parameters for the executive function-activating curve stay constant as the evaluation of the shuffled test set proceeds, the magnitude of the strength parameter for the prepotent impulse curve increases at a constant rate k each time an MNIST image containing the same handwritten digit as the previously encountered MNIST image is presented but resets when an MNIST image containing a different digit is encountered. Thus, for the nth MNIST image encountered (where d is the digit contained in the image:

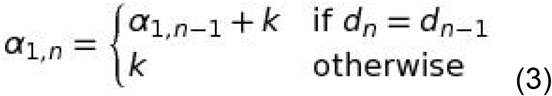

A prepotent response curve is generated by subtracting the value of the EF curve from the value of the PI curve at each point. The difference between the maximum values at each curve’s peak is used to calculate the prepotent response probability (PR) for each MNIST image:

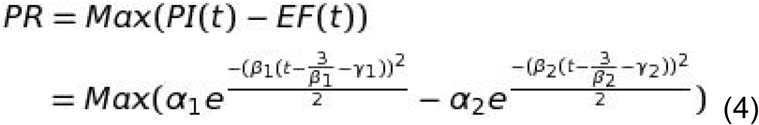

The PR value is then compared to a randomly generated value between 0 & 1 (RV). If PR < RV, then the response given by the virtual “subject” is determined by using the current MNIST image as an input for the convolutional neural network of the “subject’s” working memory representation. Otherwise, the response given by the virtual “subject” is simply a repeat of the previously given response.

The purpose of the prepotent impulsivity experiment was to determine whether this model could produce a significant difference in the performance of the working memory function. All calculations were performed with Wolfram Mathematica v11.1, and data visualizations (bar graphs) were produced with R v3.6.. In this experiment two groups of convolutional neural networks (n = 6 x networks per group) were generated with the training procedures described earlier to produce a working memory deficient group and a “healthy” working memory group. Each group’s baseline handwritten digit recognition accuracy was first evaluated on the MNIST test set. Both groups were then evaluated on six trials of shuffled test sets with the addition of the prepotent impulsivity function (all group “subjects” were evaluated with the same shuffled test set in each trial). The following set of parameters was used: k = 0.2, *β*_1_ = 0.75, *α*_2_ = 0.5, *β*_2_ = 0.25, and *γ*_1_ = *γ*_2_ = 0. These values were chosen to produce PI curves that peak rapidly and EF curves rise more slowly, similar to the relative speeds of processing for these mental activities as described in the race model. Data was tested for normality with the Shapiro-Wilk test, and all data except for the results of one “deficient” working memory neural network in the experimental condition were found to be normally distributed. Therefore, both the difference in handwritten digit recognition performance between groups and the differences in handwritten digit recognition performance within each group with the addition of the prepotent impulsivity function were tested for significance with the t-test.

Results of this experiment are shown in Figure 4. After training for MNIST handwritten digit recognition, the “healthy” working memory group achieved a mean accuracy of 95.2% (SD 1.1%) while the “deficient” working memory group achieved a mean accuracy of 55.8% (SD 2.5%) on the MNIST test set. With the addition of the prepotent impulsivity function the “healthy” working memory group achieved a group mean accuracy of 78.0% (SD 0.9%) while the “deficient” working memory group achieved a group mean accuracy of 46.5% (SD 1.9%) across all trials. Both the baseline difference in performance between the two working memory groups, and the differences in performance within each group resulting from the addition of the prepotent impulsivity function were significant. These results indicate that a statistically significant difference in handwritten digit recognition performance could be produced in both the “healthy” and “deficient” working memory groups with the addition of the prepotent impulsivity function.

**Figure 4.**
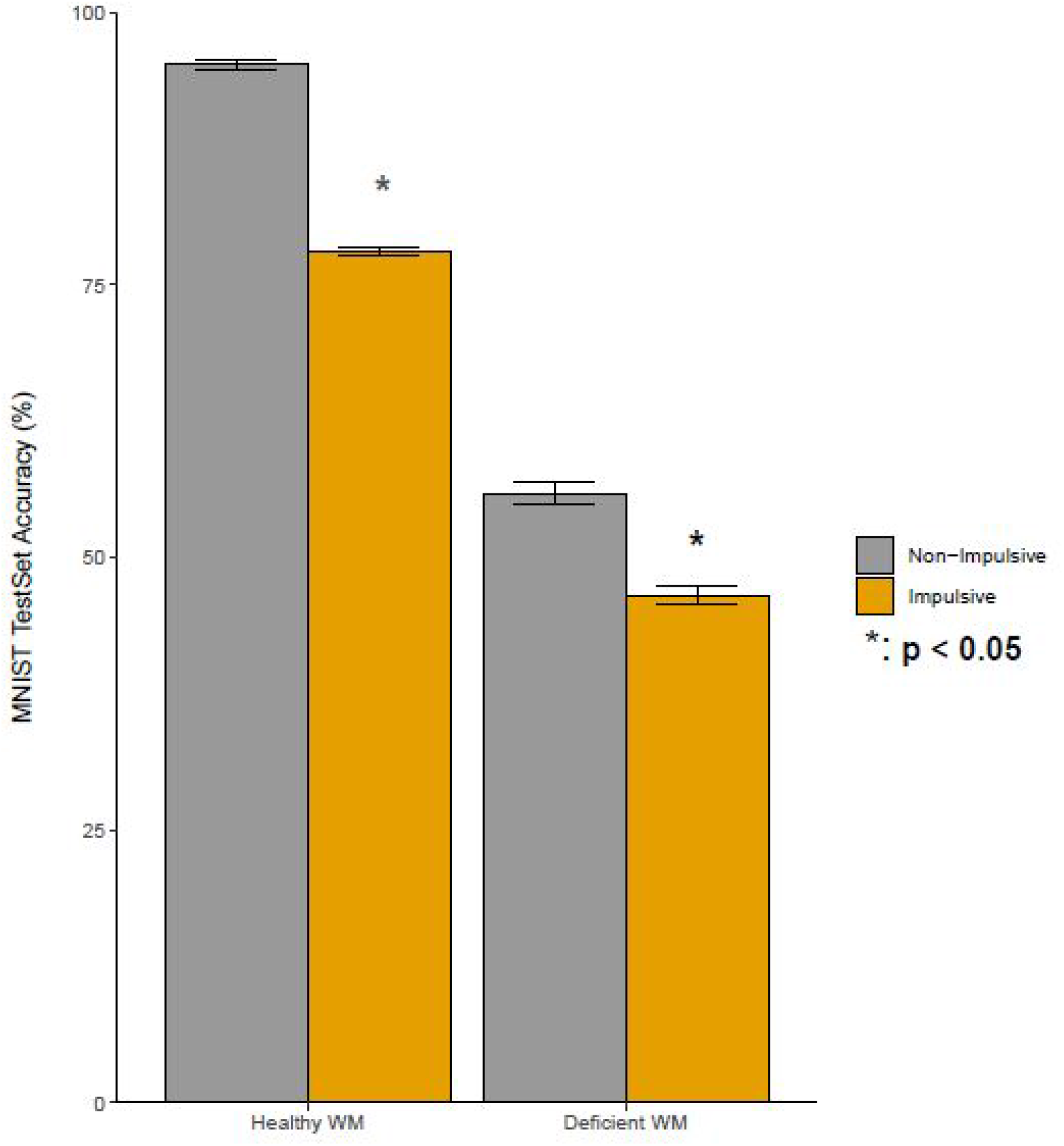
Effects of Prepotent Impulsivity Function on Group Mean Handwritten Digit Recognition Accuracy of CNNs on Shuffled MNIST TestSet (n = 6 × CNNs per Working Memory Group)

## Discussion

In summary, the beginnings of a computational model to generate personalized executive function training activity regimens for children with ADHD was developed in this work. This model utilizes a virtual “subject” constructed from a combination of core executive functions that will complete virtual executive function training activities designed to resemble executive function training activities completed by children with ADHD in a therapeutic video game intervention. The model described here utilizes convolutional neural networks that identify handwritten digits in the MNIST dataset as a representation for working memory, and includes a prepotent impulsivity function that interferes with the use of this working memory representation. Differences in working memory performance were produced by varying the number of MNIST images used for training the convolutional neural networks, and it was shown that the addition of the prepotent impulsivity function could provide an additional statistically significant effect on the performance of working memory.

Working memory is defined as the ability to hold information in consciousness (short-term memory) and manipulate it^18^. This definition leads to the rationalization for the use of image identification by convolutional neural networks as a model for working memory. In the current implementation, the input of an MNIST image file into the first layer of the network is proposed to represent the holding of information in consciousness component of working memory, and the subsequent processing of this input data by the internal network layers is proposed to represent the manipulation component of working memory. While the training processes of ANNs are of greater interest than their functional processes in the context of this work, consideration of functional modifications to make this working memory representation a more accurate representation of working memory function in humans may still be of use to improve the translational capacity of this model to human subjects.

The input of data into the network as the short-term memory component of working memory admittedly is not a strong representation of this mental ability for two reasons. First, the data provided as input to the convolutional neural network is the actual MNIST example data itself, so there is no recall mechanism to produce an internal representation of this MNIST example data in its absence. Second, there is no holding mechanism to maintain this internal representation of the data over time to allow its manipulation. Thus, the working memory function could be improved through the addition of data recall and holding mechanisms to the convolutional neural network. The recall mechanism would operate prior to the input of data to the convolutional neural network, while the holding mechanism would operate simultaneously with the processing of the input data in each hidden layer of the convolutional neural network. These modifications could potentially improve the resemblance of this working memory function to the function of working memory in biological neural networks by including a simplified representation of the biological short-term memory component while maintaining a manageable level of computational complexity.

Although several neural network models have been proposed to represent the short-term memory component of working memory^44^, a degree of redundancy may exist in the actual biological neural networks that are responsible for this executive function so no single computational process is responsible for biological working memory in all cases_45_. Two examples of artificial neural network models for modeling biological working memory include cell assemblies and synfire chains. Cell assemblies^46^ consist of strongly interconnected groups of neurons in a Hopfield model^47^ that maintain a persistent excitation pattern over time through their mutual activation. Here, the outputs of a given group of neurons at one instance in time is used as the input for this same group of neurons in the next instance to produce a recurrent firing pattern. The excitation pattern maintained by cell assemblies is a representation of a piece of information held in short-term memory, and this model relies on the use of the leaky-integrator differential equation to model the temporal dynamics of input currents and firing rates of individual neurons. Synfire chains^48^ also rely on the leaky-integrator differential equation to model their temporal dynamics, but use a feedforward neural network model rather than a recurrent Hopfield model. While the standard feedforward neural networks have the individual neurons in each layer firing at varying rates, individual groups of neurons/layers in synfire chains all fire simultaneously in time, and produce a persistent chain of spikes across multiple layers that represent the piece of information held in short-term memory. These models have been developed from the biophysical properties of individual neurons’ dynamic function, and therefore attempt to capture a level of complexity that is too great for the purposes of this project at this time. Nevertheless, much of this complexity may be obviated with the selection and design of simpler models that ignore these temporal dynamics of neural function while still representing similar functions.

To represent the recall mechanism of short-term memory, a generative learning algorithm^49^ could be added to the current working memory model. In contrast to discriminative models (such as convolutional neural networks) that transform input data into a classification category, generative models can receive a classification category as an input and reconstruct data examples that correspond to the input category. This type of network could be trained to produce an MNIST image as an output when a digit in the range 0-9 is provided as an input. By modifying the training parameters of the generative model, the quality of the MNIST images produced can be varied. Hence, a properly trained generative network with an effective recall would reproduce high-quality MNIST images as outputs when provided with digit labels as inputs, whereas a poorly trained generative network with an ineffective recall would produce poor-quality MNIST images as outputs. These reproduced versions of MNIST images would then be provided as inputs to the manipulation function represented by convolutional neural networks in the current implementation of the working memory model. The high-quality MNIST images produced by well-trained generative networks with effective recall would enable more effective performance of the manipulation function represented by convolutional neural networks and result in a higher probability of a correct handwritten digit identification and more effective use of working memory as a whole. In contrast, the low-quality MNIST images produced by poorly-trained generative networks with ineffective recall would impede the performance of the manipulation function represented by convolutional neural networks and result in a lower probability of a correct handwritten digit identification and less effective use of working memory as a whole.

One potential approach to represent the holding mechanism of short-term memory is to interfere with the passage of information between individual layers of the convolutional neural network during the operation of the manipulation function. Although the original input data is transformed with each passage from layer to layer, it can still be considered a modified representation of the original information and therefore the act of transferring the output from one layer as an input to the next layer could be considered analogous to holding the information being manipulated. Within this context, a dysfunctional holding mechanism could be represented with an ineffective transfer of information between layers (by randomly corrupting the data values before they are input into a layer, for example) and an effective holding mechanism could be represented by an unimpeded transfer of information between layers.

The design of the impulsivity function was influenced by both the race model^34^ and the passive-dissipation hypothesis^35^ for the control of impulsive behavior. These theories have been used to explain the results observed in the stop-signal task, which is a behavioral task requiring the use of inhibitory control. Both of these theories describe impulsive responses and their inhibition in terms of mental processes that grow in strength over time and compete to reach a threshold value that activates a behavioral response. These theories differ in the mechanism of impulsive behavior prevention, as the passive-dissipation hypothesis includes an active role for inhibitory control (described below). The impulsivity function in the current project was designed to represent the mental processes that compete to produce behavioral responses as described in these theories. However, the role of the impulsivity function at this time is the generation of impulsive behaviors rather than their inhibition, although the design of an inhibitory control component that works as described in the passive-dissipation hypothesis remains to be addressed in future work.

The race model has been used to describe the mental processing of prepotent responses and their inhibition in the stop signal task^34^. Prepotent responses are a category of behavioral responses to stimuli for which immediate reinforcement is presently available or has been available in the past, and were identified as a target for the behavioral inhibition executive function in Barkley’s influential executive function theory for ADHD^16^. Prepotent responses can be overlearned behaviors, occur impulsively without reflection, and can oftentimes conflict with long-term goals. All individual trials in the stop signal task contain a “go” signal that requires the experimental subject to select a response. A subset of these trials also include the presentation of a “stop” signal at varying time intervals after the “go” signal has been presented, and require the experimental subject to withhold the selection of a response. The presentation of the “stop” signal after the “go” signal is significant in that the mental processing for responding to the “go” signal is already active when the “stop” signal is presented. The race model proposes that these processes are independent and compete to produce a behavioral response, and the resulting behavioral response is activated by the process that reaches a given threshold more rapidly in time.

The passive-dissipation hypothesis^35^ builds on the race model by allowing the prepotent response to be overcome by the use of the executive function of inhibition to create a delay in the decision to respond. This delay allows the slower correct response to continue growing in strength to reach the response threshold as the prepotent response dissipates. While the use of the executive function inhibition is not included in this project, the impulsivity function was designed to allow for the use of inhibition to create a delay in the decision to respond. When this delay occurs the slower mental process for an appropriate response grows in strength and exerts a greater influence on the decision-making process.

Similar to the impulsivity function, both the race model and passive-dissipation hypothesis describe impulsive behaviors and their inhibition in terms of competing mental processes. However, while the race model and passive-dissipation hypothesis define this competition as a race in time between mental processes to reach a given threshold value, the impulsivity function defines this competition in terms of the relative magnitudes of the outputs of the mental processes. In other words, the race model and passive-dissipation hypothesis describe the resulting behavioral response as an outcome of only one of the mental processes, while the impulsivity function defines the behavioral response as an outcome of an interaction between both mental processes. In the race model the winning process is simply the process which reaches the threshold first, while in the passive-dissipation hypothesis the winning process is the one which reaches the threshold when the decision to act is made. In contrast, the impulsivity function implements the resulting behavioral response as the outcome of an interaction between both of the mental processes by subtracting the strengths of the outputs of the mental processes.

This initial implementation of the impulsivity function was designed to create impulsive responses to interfere with the operation of the working memory function in the absence of any influence from behavioral inhibition. Thus, the current version of the impulsivity function simply defines the decision point as the time when the PR process curve reaches its peak value, which is when the PR process exerts its maximum effect on the behavioral response and the EF process exerts a relatively minor effect. However, a behavioral inhibition function may be added in future implementations of the impulsivity function to create a delay of the decision point to allow for the PR process to weaken as the EF process strengthens, leading to a greater opportunity for the application of a productive behavioral response as reflected in a decreased PRP value.

Two alternative computational methods that have been used for modeling inhibition are the negative bias weight learning mechanism and the drift-diffusion model. The negative bias weight learning mechanism^50^ was applied within the context of a dynamic gating neural network model for a task-switching activity similar to the Wisconsin card sorting task, which is a psychological test used to measure cognitive flexibility^51^. The negative bias weight learning mechanism inhibits responses by rapidly decreasing the weight values for individual units in the neural network to make them inactive. These weights are then allowed to gradually increase to zero as the task continues to lift the inhibition. The negative bias weight learning mechanism operates within the context of the continuous training of a neural network as it performs a given task. In contrast, the impulsivity function operates as an external precursor to the CNNs in this model, and therefore the negative bias weight learning mechanism would not be compatible with this current approach.

A potential alternative method for the computational modeling of behavioral inhibition processes is the drift diffusion model^52^. The drift diffusion model has garnered interest in computational psychiatry to model decision-making processes^53^, and the parameters of this model have been linked to neural functions^54–57^. It has been useful for modeling simple one- or two-choice decisions^54^, where the selection of a decision is determined by one or more stochastic drift processes that move towards boundaries representing the available decisions. The stochasticity in the model is incorporated through the addition of random noise to the movement of the drift processes as they travel towards a boundary. When the movement of the drift process reaches one of the boundaries, a decision to complete the action represented by that boundary is made.

In contrast to the negative bias weight learning mechanism, the drift-diffusion model offers a viable alternative for the representation of impulsive behaviors and their inhibition currently within the context of the prepotent impulsivity function developed here. While the drift-diffusion model has a stronger record of evidence for its use, to our knowledge its application has been limited to the modeling of decision-making in simple one- or two-choice decision-making tasks. A complex gaming environment such as Minecraft, however, presents dynamic decision-making scenarios where more choices may be available at any given moment and choices are continuously removed or added as circumstances change. It is foreseeable that higher complexity environments could present decision-making scenarios that require alternative criteria for inhibitory decision-making that extend beyond the time-based processing of alternatives by the drift-diffusion model. Therefore, the use of multiple inhibitory decision-making models may be advantageous in this project to determine whether different models may work more effectively in different varieties of decision-making scenarios.

In conclusion, a unique computational representation of working memory and the mental processes that generate impulsive behavior was developed in this project. This model incorporates the training process of artificial neural networks as a potential method for predicting personalized executive function training schedules in children with ADHD. As this model is still in its preliminary stages, it will be necessary to incorporate representations of other executive functions and develop virtual executive function training activities that rely on these representations for their completion. Finally, the utility of this approach to guide the selection of therapeutic interventions remains to be determined.

## Acknowledgements

The project described was supported in part by the National Center for Research Resources and the National Center for Advancing Translational Sciences, National Institutes of Health, through Grant UL1TR001998, and by NSF ACI-1053575 allocation number BIO170011. The content is solely the responsibility of the authors and does not necessarily represent the official views of the NIH or NSF.

